# The modulation of acute stress on Model-Free and Model-Based reinforcement learning in Gambling Disorder

**DOI:** 10.1101/2022.05.05.490735

**Authors:** Florent Wyckmans, Nilosmita Banerjee, Mélanie Saeremans, Otto Ross, Charles Kornreich, Laetitia Vanderijst, Damien Gruson, Vincenzo Carbone, Antoine Bechara, Tony Buchanan, Xavier Noël

## Abstract

**Background and aims:** Experiencing acute stress is common in behavioral addictions such as gambling disorder. Additionally, like most substance-induced addictions, aberrant decision-making wherein a reactive habit-induced response (conceptualized as a Model-free [MF] in reinforcement learning) suppresses a flexible goal-directed response (conceptualized as a Model-based [MB]) is also common in gambling disorder. In the current study we investigated the influence of acute stress on the balance between habitual response and the goal-directed system.

**Methods:** A sample of *N = 116* pathological gamblers (PG) and healthy controls (HC) performed an acute stress task – the Socially Evaluated Cold pressure task (SECPT) – or a control task. Self-reported stress and salivary cortisol were collected as measures of acute stress. Following the SECPT, participants performed the Two-Step Markov Task to account for the relative contribution of MB and MF strategies. Additionally, verbal working-memory and IQ measures were collected to account for their mediating effects on the orchestration between MB/MF and the impact of stress.

**Results:** Both groups had comparable baseline and stress-induced cortisol response to the SECPT. Non-stressed PG displayed lower MB learning than HC. MANOVA and regression analyses showed a deleterious effect of stress-induced cortisol response on the orchestration between MB and MF learning in HC but not in PG. Neither working memory nor IQ mediated these effects.

**Discussion and Conclusions:** Despite normal cortisol response to stress, we found an abnormal pattern of modulation of stress on the orchestration between MB and MF learning among PG.

## INTRODUCTION

Gambling is a ubiquitous recreational activity worldwide. Despite its recreational facet, for a small yet significant proportion of gamblers (estimated around 0.1% to 5.8% worldwide), tenacious engagement in gambling results in the development of Gambling Disorder (GD), a persistent, excessive and uncontrollable urge towards gambling despite the looming negative consequences of this excessive engagement (APA, 2013). Two key aspects that characterize GD are: 1) gambling to escape stress (Buchanan et al., 2020) and 2) making decisions that are informed by habit-induced, reactive responses instead of using a deliberative, goal-directed flexible approach (Everitt & Robbins, 2005). While these characteristic features of GD have been studied in isolation, to our knowledge a systematic investigation of a possible interaction between the experience of acute stress and its consequent impact on habit versus flexible decision-making has not been undertaken in GD. The aim of this study is to investigate the existence of this possible interaction. More specifically, the current study aims to examine stress-induced modulation of the arbitration between habit-induced, highly reactive decisions versus deliberative, goal-directed flexible decisions.

Stress represents a major accelerating factor in the development of addictive disorders (Enoch, 2011), including GD (Biback & Zack, 2015; Buchanan et al., 2020; Oakes et al., 2019). Stress has a paradoxical impact in GD. On the one hand recreational elements of gambling activities and the rewarding experience of gambling engagements are leveraged by gamblers to cope with stressful life events (Edgerton et al., 2018; Weinstock et al., 2008), on the other hand consistent gambling engagement in itself can be a distressing experience (APA, 2013) making gambling a potential stressor (Russell et al., 2021). From a neurocognitive perspective, recurrent stress exposure can cause profound changes within neural systems involved in decision-making (Wirz et al., 2018). Particularly, recurrent exposure to acute stress may increase one’s tendency to behave in an inflexible habitual way (Schwabe & Wolf, 2009, 2010, 2011; Seehagen et al., 2015).

Clinically, inflexible and habitual responses (as opposed to deliberative and goal-directed responses) inform the decisions undertaken in addictive states like GD (c.f., Wyckmans et al., 2019), fueling the persistence of addictive behaviors like gambling despite the possibility of recurring negative consequences (e.g., financial losses; Everitt & Robbins, 2005). According to reinforcement learning (RL) theory, such habitual, inflexible responses and the contrary goal-directed behaviors arise from two parallel learning systems, Model-Based (MB) and Model-Free (MF), which differ in their way of updating choice-value during instrumental learning (Daw et al., 2005, 2011). According to RL theory, when facing a decision, the MB-system relies on an internal model of the contingencies between actions and consequences, which informs the computations of the expected values for each candidate decision (Daw, 2018). Thus, MB strategies underlie goal-directed behavior, as it enables flexible adaptation to the environment and prospective update of action value in view of long-term goals (Dolan & Dayan, 2013). Conversely, the MF-system bypasses these laborious simulations and directly computes the action-value based on past reward history (Rummery & Niranjan, 1994; Sutton & Barto, 1998). Therefore, the MF-system is associated with habitual behavior as it depends solely on previously learned associations (Dolan & Dayan, 2013). Despite the advantage gained by the lowered cognitive burden in MF decision-making strategies, it comes at the cost of hampered flexibility: if the contingencies between action and consequences change, the stored action values become obsolete and lead to outdated choices (Daw, 2018). Thus, under RL theory, addictive behaviors (which occur from excessive reliance on habitual, reflexive responses coupled with diminished reliance on deliberative goal-directed systems), arise from the asymmetrical dynamics between the MF-MB systems wherein there is an exacerbated MF-system involvement coupled with a suppressed MB-system involvement underlying the persistence of the addictive behaviour at hand (e.g., Groman et al., 2019). In fact, recent research undertaken in the field of substance-induced addictions (e.g., binge drinking; Doñamayor et al., 2018) highlights that the presence of such an imbalance between the MF and MB systems acts as a key factor underlying the perpetuation of the addictive behaviours. Crucially, recent evidence also suggests that a preclinical existence of such imbalance could predispose an individual to develop addictive behaviours (e.g., Chen et al., 2021). Additionally, evidence from recent research has also replicated these findings with PG (i.e., in the behavioral addiction of GD; Wyckmans et al., 2019). Wyckmans and colleagues (2019) demonstrated a diminished MB-system involvement coupled with an heightened MF-system involvement in GD in the Two-step Markov task (Daw et al., 2011) that systematically accounts for the MF/MB strategies in decision-making.

In terms of the impact of stress on MF/MB-systems, studies indicate that acute stress has a deleterious effect on MB learning (Otto, Raio, et al., 2013; Radenbach et al., 2015) which occurs due to the hampered functioning of the amygdala-dorsal striatum connectivity and prefrontal cortex, resulting in diminished executive processes (Wirz et al., 2018). Conversely, stress-induced cortisol release enhances MF learning (Park et al., 2017) by increasing the firing rate of dopaminergic neurons in the dorsal striatum (Anstrom & Woodward, 2005), a region involved in habit formation (Everitt & Robbins, 2005, 2016) such as those observed in addictive states. Considering the above evidence, it seems plausible to reason that repeated exposure to acute stress during gambling might present a risk factor for the development of GD by favoring MF learning, thus rendering the resulting behavior resistant to its negative consequences. Therefore, acute stress experienced from gambling (Russell et al., 2021), the increased reliance on MF-systems among PG (Wyckmans et al., 2019), and the amplification of MF-systems as a function of increased acute stress(Park et al., 2017), calls for a need to systematically investigate the effects of acute stress on the MF-system among PG.

To pursue this line of investigation and understand how acute stress modulates the arbitration between MF/MB-systems, computational modelling approaches that augment the previously used experimental inferential analyses approaches are needed (e.g., Wyckmans et al., 2019). Traditionally, MF/MB-systems are studied using the Two-step Markov task (RL-task; Daw et al., 2011). Experimental approaches which focus on inferential analyses of the choice behavior data such as reaction times and state-transition probabilities from the Two-Step Markov Task do not provide insights about the latent learning processes in decision-making that arise from the implementation of MF or MB strategies. Discerning these latent psychological variables is crucial to account for the impact of acute stress on the relative weighing between MF/MB systems. Thus, fitting behavioral choice data from the Two-Step Markov Task to a Hybrid-RL 7 parameter model (Daw et al., 2011), wherein each parameter will allow us to systematically estimate the learning processes involved during decision-making (Gillan et al., 2015; Mollick & Kober, 2020), would appropriately discern the impact of acute stress on the dynamics between MF and MB.

The aim of the current study is to systematically investigate the impact of acute stress on the exacerbation of the MF-system among PG. We aim to investigate this by fitting a Hybrid-RL 7-parameter computational model to the behavioral data obtained from Two-step Markov task, a novel approach which remains unexplored in this line of research. Acute stress will be induced with the Socially Evaluated Cold Pressor Task (SECPT) that requires participants to immerse one arm into cold water while being socially observed and videorecorded, a method that robustly increases laboratory stress reactivity (Schwabe et al., 2008), including among PG (Grant & Chamberlain, 2019). Based on previous findings we hypothesized that PG will repeat previously rewarded choices without accounting for the task structure (Wyckmans et al., 2019). Thus, we expected to observe a general bias towards MF learning among PG as compared to healthy controls. Furthermore, we hypothesized that stress would further increase the activity of the MF-system and suppress the MB-system among stressed PG as opposed to stressed controls. Finally, we also accounted for verbal working memory and abstract reasoning on MB/MF to account for their mediating effects on the acute stress and MF/MB interaction.

## METHODS

### Participants

Seventy-three pathological gamblers (PG) with a score of 6 or higher on the South Oaks Gambling Screen (Lesieur & Blume, 1987) and 75 healthy controls (HC) were recruited. Only male participants (to avoid the confounding effect of menstrual cycle on cortisol response to stress; Montero-López et al., 2018) over 18 and without a history of neurological or psychiatric diseases were included. We excluded five participants (two PG and three HC) due to corrupted cortisol samples, as well as 27 participants (13 PG and 14 HC) who did not meet the success criteria for the RL-Task (see below), yielding a final sample of 116 participants (58 PG and 58 HC).

### Procedure

The entire procedure lasted two hours (figure 1). Remuneration was set at 30€, with up to 10€ extra depending on individual task performance. Participants first underwent a semi-structured interview assessing demographic characteristics and gambling habits (if applicable). Fluid intelligence was assessed with the nine-item forms of the Raven’s Standard Progressive Matrices Test (see supplementary material, Bilker et al., 2012), followed by a working memory task, the Operation Span Task (OSPAN; see supplementary material, Unsworth et al., 2005) and the RL-task instructions. Before completing the RL-task, half the participants underwent the SECPT (Schwabe et al., 2008), during which they immersed their forearm in a basin of cold water (3°c) for three minutes while being closely observed by an experimenter and videorecorded. The other half underwent the control procedure (Warm Pressor Task; WPT), during which the water was at ambient temperature and no camera or observation was used. In total, 60 and 56 participants underwent the SECPT (30 HC and 30 PG) and the WPT (28 HC and 28 PG), respectively. Participants finished the session by filling out clinical questionnaires (see supplementary material).

**Figure 1.**
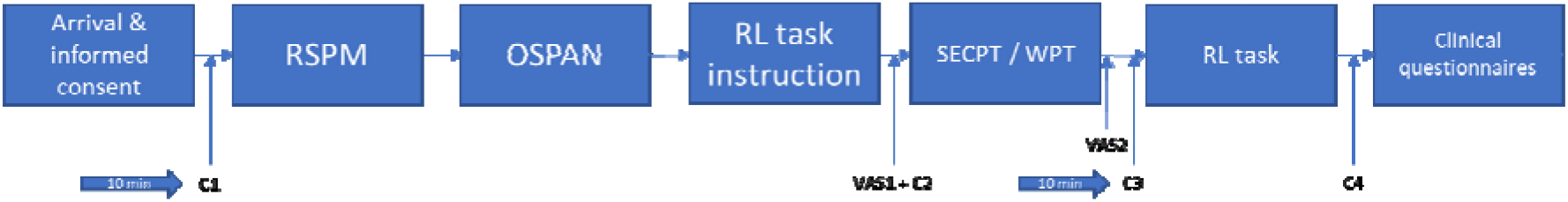
Experimental protocol. After filling out the informed consent form, participants underwent the Raven’s Standard Progressive Matrices Test (RSPM) and OSPAN task. The first cortisol measurement (C1) was collected 10 minutes after the arrival. The second (C2) was collected right after the dual-step Markov Task (RL task) instruction along with the first self-reported evaluation (VAS1). The second VAS was taken right after the stress induction (SECPT) / control (WPT) procedure, while the third cortisol measurement (C3) was collected 10 minutes after the procedure. The fourth measurement (C4) was collected right after the Two-Step Markov Task (RL task). Participants finished by filling out several questionnaires.

### Measures

#### Stress response assessment

Participants rated their desire to gamble, feeling of stress, and pain on visual analog scales (VAS; ranging from 0 to 10) before and after the stress induction procedure. Objective stress response measures were taken in the form of salivary cortisol levels, collected 10 minutes after arrival (C1), immediately following the RL-task instructions (C2), 10 minutes post-stress induction (C3), and after the RL-task (C4). C1 and C3 were delayed by 10 minutes as cortisol levels are expected to peak 10 minutes after stress onset (McRae et al., 2006). Salivary cortisol analysis procedure is described in the supplementary material.

#### Two-Step Markov Task

All participants received extensive instructions for the Two-Step Markov Task (RL-task; Daw et al., 2011; Otto, Raio, et al., 2013; Wyckmans et al., 2019) and completed 10 practice trials at the end of which they answered three instruction-related questions. The instructions were reiterated in case of incorrect responses. Participants subsequently performed 200 trials of the RL-task (figure 2A). During the first step, they were required to choose between two fractal images displayed alongside each other on a black background. Each image commonly (70%) or rarely (30%) led to one of the two second-step stages (figure 2C). Participants were informed that these probabilities would stay fixed. During the second step, they had to choose between two fractal images displayed alongside each other on a colored background (green or blue, depending on their first-step choice). Reward probabilities for each fractal image varied in function of background color. Participants were informed that these second-step probabilities would slowly fluctuate as the task progressed (following a Gaussian Random Walk with boundaries fixed at 0.25 and 0.75; SD = 0.025; figure 2B) to ensure continual exploration. After the second step, a 1s feedback slide was displayed to indicate whether the trial was rewarded (a 20c coin) or not (“0”). Participants had 3s to indicate their preference by pressing the letter “E” (left image) or “I” (right image) on an AZERTY keyboard. Inter-stage and inter-trial intervals lasted 1s each. Before data analyses, we automatically excluded participants who failed to answer within 3s more than 20 times, who picked the same first-step choice in 95% of the trials, or who repeated previously rewarded second-step responses at a rate lower than 50%. MB and MF algorithms predict different observable choice patterns during subsequent 2-step trials (figure 2D). Under MF strategy, decisions will be reinforced solely depending on the outcome of the previous trial. Conversely, choice-value will update depending on the interaction between previous outcomes and transitions under MB strategy.

**Figure 2.**
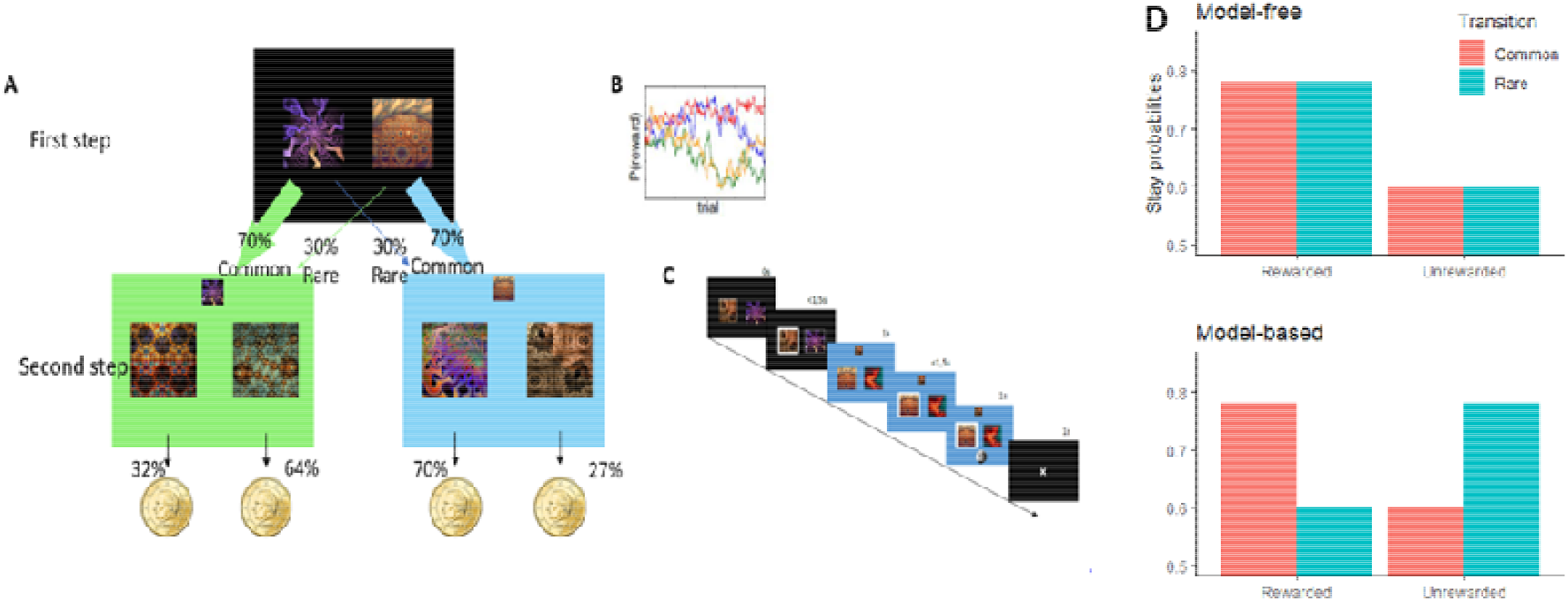
A. Two-step decision task. (First step) Participants chose between the two images, leading preferentially to a green or a blue screen, according to fixed probabilities. (Second step) Subjects chose between the two images linked to probabilities to win money. Those probabilities slowly changed with time and differed according to the background color. B. Second step’s changes in the probability of reward. C. Trial’s design. D. Theoretical decision pattern according to a pure MF strategy and to a pure MB strategy.

### Statistical analyses

Data cleaning and statistical analyses were performed using RStudio (v1.4.1103) and SPSS (v28). The sample size was estimated based on previous studies (Sebold et al., 2014; Voon et al., 2015). Accordingly, we aimed at finding a medium-sized difference between HC and PG with 80% power and 95% confidence, which required 64 participants per group.

Pre- and post-procedure cortisol concentrations were computed by averaging the first two and last two measures, respectively. Results were log-transformed because of the skewed nature of their distribution (Petzold et al., 2010). The influence of the SECPT/WPT procedures on cortisol concentrations and subjective ratings were assessed through repeated measures MANOVA. Our total sample was split into two groups depending on their cortisol response. Participants with a cortisol increase equal to or higher than 1.5 nmol/l (N = 25) were considered cortisol responders (Miller et al., 2013).

We fitted choice behavior on the RL-task to a 7-parameter hybrid reinforcement learning algorithm (Daw et al., 2011) with the hBayesDM package (Ahn et al., 2017). Four MCMC chains of 6000 samples each were run, with the first 3000 samples as warm-up (see supplementary material). We mainly focused our analyses on the *ω*-parameter, which indicates the relative contribution of MB learning over MF learning during task completion. A between-subject MANOVA was used to assess the influence of stress group (Stressed vs. Non-stressed), diagnostic group (PG vs. HC), and their interaction on these 7 parameters. Multiple regression was performed to assess the continuous effect of cortisol increase, diagnostic group, and their interaction on the *ω*-parameter.

Normality was assessed by dividing the skewness and excess kurtosis by their respective standard deviation. Distributions with both scores [-3.29; 3.29] were considered normal (Kim, 2013). The continuous variables were standardized before each regression. Demographic and clinical variables from both groups were compared with Mann-Whitney U or Welch t-tests. To avoid the confounding effect of outliers in regressions, scores that differed over 3 times the MAD from the median (Leys et al., 2013) were removed before each regression.

### Ethics

Participants were recruited through advertisement and gave written informed consent to be part of the experiment. The experiment was approved by the C.H.U. Brugmann Ethics Committee (n° OM 026) and performed according to the Declaration of Helsinki.

## RESULTS

### Sample characteristics

Our final sample consisted of 116 participants, 58 HC, and 58 PG. Both groups were matched for age, gender (only males), and education. Table 1 depicts the demographic and clinical variables of PG and HC as well as between-group comparisons and Cronbach’s alpha for each questionnaire. PG displayed significantly lower scores on working memory and fluid intelligence assessments than HC, as well as significantly more psychiatric comorbidities, state-anxiety, depressive symptoms, and reward sensitivity, which were included as covariates in separate analyses (see supplementary material).

**Table 1.** Descriptive scores of each clinical variable and their between-subject difference. Clinical variables assessed alcohol use disorder symptoms (AUDIT), gambling disorder symptoms (SOGS), number of DSM items (DSM), working memory performances (OSPAN), Raven score, psychiatric comorbidities (SCL90R), depressive symptoms (BDI), positive affects (PANAS pos), negative affects (PANAS neg), state-anxiety (STAI-YA), trait-anxiety (STAI-YB), adverse live events (SRRS), sensibility to punishment (PS), sensibility to reward (RS), routine tendencies (CoH R), automatism tendencies (CoH A), negative urgency (NU), positive urgency (PU), lack of premeditation (LoPr), lack of perseverance (LoPe), and sensation seeking (SS). Groups are compared with Welsh’s t-tests or Mann-Whitney U according to the distribution of their scores. Cronbach’s alphas are displayed when applicable. Significant group differences are displayed in bold.

| Variable | PG (n = 58) | HC (n = 58) | Test | Alpha |
| --- | --- | --- | --- | --- |
| Age | 29.22 (8.39) \| Mdn = 28 | 31.16 (9.86) \| Mdn = 28.5 | U(116) = 1498.5, p = .31 | N/A |
| Study Level | 13.88 (2.85) \| Mdn = 15 | 14.66 (2.4) \| Mdn = 15 | U(115) = 1400, p = .14 | N/A |
| AUDIT | 8.87 (7.75) \| Mdn = 8 | 6.34 (4.49) \| Mdn = 5 | U(114) = 1748, p = .21 | .86 |
| SOGS | **9.05 (2.63) \| Mdn = 8.5** | **0.36 (1.1) \| Mdn = 0** | **U(79) = 1624, p < .001** | .86 |
| DSM | **3.97 (2.62) \| Mdn = 4** | **0.11 (0.32) \| Mdn = 0** | **U(90) = 1881.5, p < .001** | .88 |
| Smoker (Y\|N) | 22 \| 36 | 19 \| 39 | X (1) = 0.34, p = .56 | N/A |
| OSPAN | **0.76 (0.13) \| Mdn = 0.77** | **0.8 (0.16) \| Mdn = 0.85** | **U(116) = 1207, p = .01** | N/A |
| Raven | **4.5 (1.88) \| Mdn = 5** | **5.33 (1.94) \| Mdn = 6** | **U(116) = 1214.5, p = .01** | N/A |
| SCL90R | **55.67 (49.01) \| Mdn = 44** | **35.1 (32.54) \| Mdn = 30.5** | **U(116) = 2119.5, p = .02** | .97 |
| BDI | **6.28 (5.94) \| Mdn = 5** | **3.71 (4.12) \| Mdn = 2** | **U(116) = 2090.5, p = .02** | .88 |
| PANAS pos | 34.31 (7.8) \| Mdn = 34 | 36.21 (6.27) \| Mdn = 37 | t(108.98) = 1.44, p = .15 | .85 |
| PANAS neg | 20.6 (8.25) \| Mdn = 18 | 18.1 (6.85) \| Mdn = 16.5 | U(116) = 1970, p = .11 | .89 |
| STAI-YA | **36.54 (10.95) \| Mdn = 35** | **31.03 (8.83) \| Mdn = 29** | **t(107.39) = 2.97, p = .004** | .90 |
| STAI-YB | 42.36 (12.3) \| Mdn = 41.5 | 39.72 (11.18) \| Mdn = 38.5 | t(110.12) = 1.2, p = .24 | .91 |
| SRRS | 244.96 (169.44) \| Mdn = 201 | 242.58 (206.81) \| Mdn = 184 | U(115) = 1692.5, p = .58 | N/A |
| PS | 41.84 (11.54) \| Mdn = 41 | 39.26 (9.46) \| Mdn = 39 | t(108.08) = 1.31, p = .19 | .90 |
| RS | **43.98 (10.95) \| Mdn = 44** | **39 (7.89) \| Mdn = 40** | **U(116) = 2127, p = .01** | .88 |
| CoH R | 52.44 (12.69) \| Mdn = 53 | 45.77 (12.05) \| Mdn = 45 | t(31.46) = 1.63, p = .11 | .86 |
| CoH A | 33.94 (12.14) \| Mdn = 34.5 | 27 (9.42) \| Mdn = 22 | t(27.28) = 1.91, p = .07 | .88 |
| NU | 8.76 (3.35) \| Mdn = 9 | 7.67 (3.2) \| Mdn = 8 | t(113.76) = 1.78, p = .08 | .85 |
| PU | 10.17 (3.01) \| Mdn = 10.5 | 9.9 (2.59) \| Mdn = 10 | t(111.48) = 0.53, p = .60 | .75 |
| LoPr | 9.12 (3.5) \| Mdn = 8 | 8.47 (3.13) \| Mdn = 8 | U(116) = 1828, p = .42 | .88 |
| LoPe | 9.24 (3.63) \| Mdn = 9 | 9.07 (3.4) \| Mdn = 8 | t(113.51) = 0.26, p = .79 | .92 |
| SS | 10.38 (3.28) \| Mdn = 11 | 9.78 (3.5) \| Mdn = 10 | t(113.51) = 0.96, p = .34 | .86 |

A repeated measures MANOVA was performed to evaluate the effect of stress induction procedure (between-subject; SECPT vs. WPT), time (within-subject; before vs. after the procedure) and their interaction on salivary cortisol concentrations, as well as self-reported stress, desire to gamble (craving), and pain measures (figure 3). The interaction effect on the combined dependent variables was significant (F(4, 111) = 7.49, p < .001; Wilks’ Λ = .73, η^2^_p_ = .28). Univariate analyses showed a significant effect of the interaction for each dependent variable (table 2), indicating a higher increase in cortisol concentration and subjective ratings among participants undergoing the SECPT than the WPT. No group effect (PG vs. HC) was found on pre- to post-procedure differences in cortisol concentration and subjective ratings (F(4, 111) = 0.96, p = .43; Wilks’ Λ = .97, η^2^_p_ = .03).

**Figure 3.**
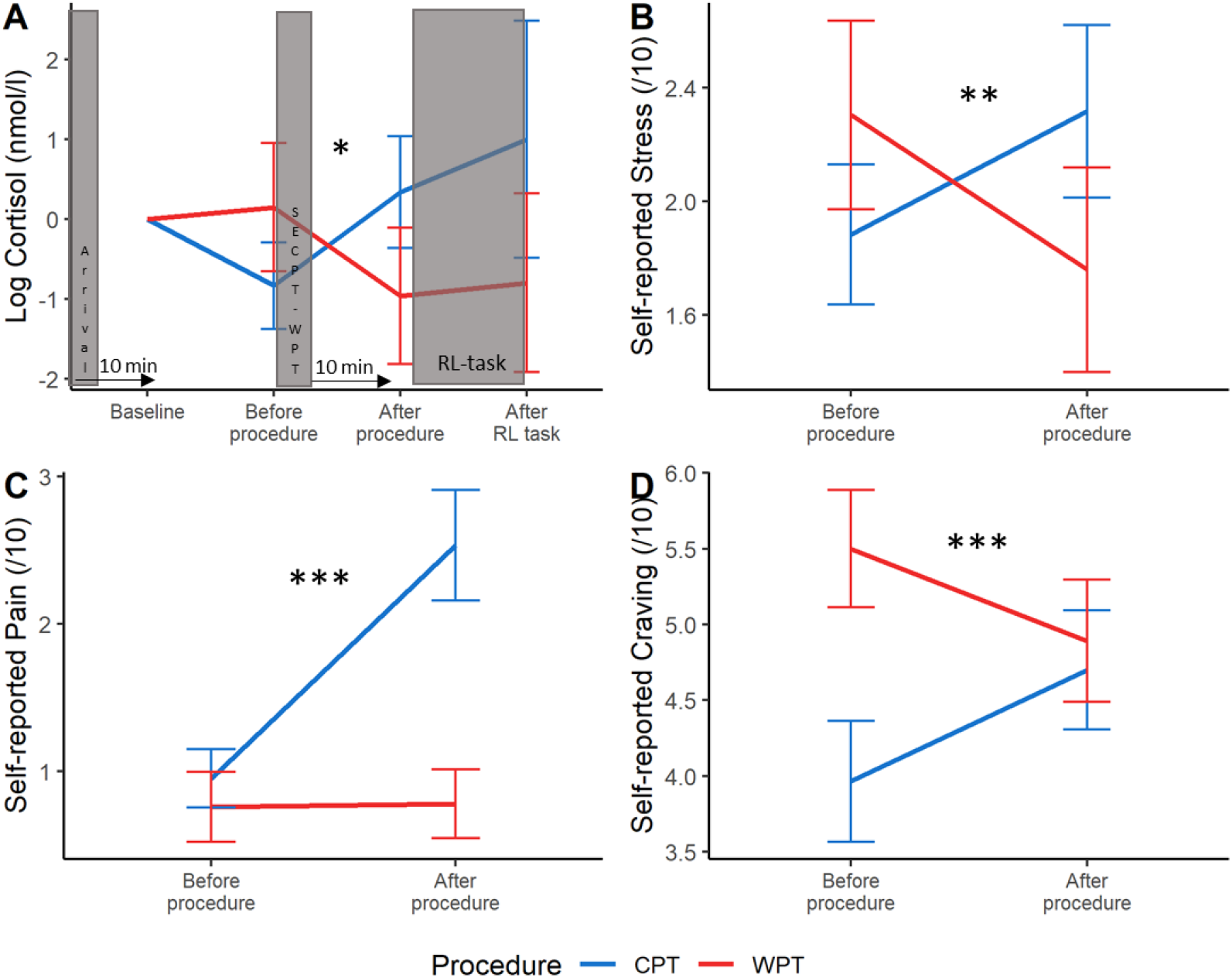
Cortisol and self-reported measures for the group that underwent the cold-pressor task (SECPT) and the Warm-Pressor Task (WPT). Graphs show mean values ± SE. P-values of the effect of interaction between time and procedure are reported * p < .05, ** p < .01, ***p < .001.

**Table 2.** Mean (sd) cortisol concentrations (not log-transformed for interpretability) and self-reported measures, before and after the pressor task. Results of the univariate interactions between time and procedure type are displayed.

|  | Cold Pressor Task (n = 60) |  | Warm Pressor Task (n = 58) |  | Univariate results |
| --- | --- | --- | --- | --- | --- |
|  | Before Procedure | After Procedure | Before Procedure | After Procedure | Time*Procedure |
| Cortisol concentration (nmol/l) | 5.94 (4.11) | 7.02 (6.70) | 7.63 (5.38) | 6.68 (6.53) | F(1,114) = 9.41,<br>p = .003, $\eta^2_p = .08$ |
| Stress (/10) | 1.88 (1.91) | 2.32 (2.35) | 2.30 (2.48) | 1.76 (2.70) | F(1,114) = 8.86,<br>p = .004, $\eta^2_p = .07$ |
| Pain (/10) | 0.95 (1.53) | 2.53 (2.90) | 0.76 (1.78) | 0.78 (1.76) | F(1,114) = 17.69,<br>p < .001, $\eta^2_p = .13$ |
| Craving (/10) | 3.32 (3.11) | 4.12 (3.20) | 4.07 (3.4) | 3.79 (3.25) | F(1,114) = 10.61,<br>p = .001, $\eta^2_p = .09$ |

We divided our sample into two groups according to their cortisol response: participants with a cortisol elevation higher than 1.5 nmol/l were considered as demonstrating a cortisol response (Miller et al., 2013). The proportion of PG and HC did not significantly differ between the responders (10 PG and 15 HC) and the non-responders (48 PG and 43 HC) groups (X^2^(1) = 1.28, p = .26). The responder group contained significantly more participants who underwent the SECPT than the WPT (X^2^(1) = 5.25, p = .02).

### Analyses of choice behavior

MB and MF algorithms predict differential probabilities of repeating the previous trial’s first step’s choice depending on its outcome and transition. These differential probabilities are illustrated in figure 4.

**Figure 4.**
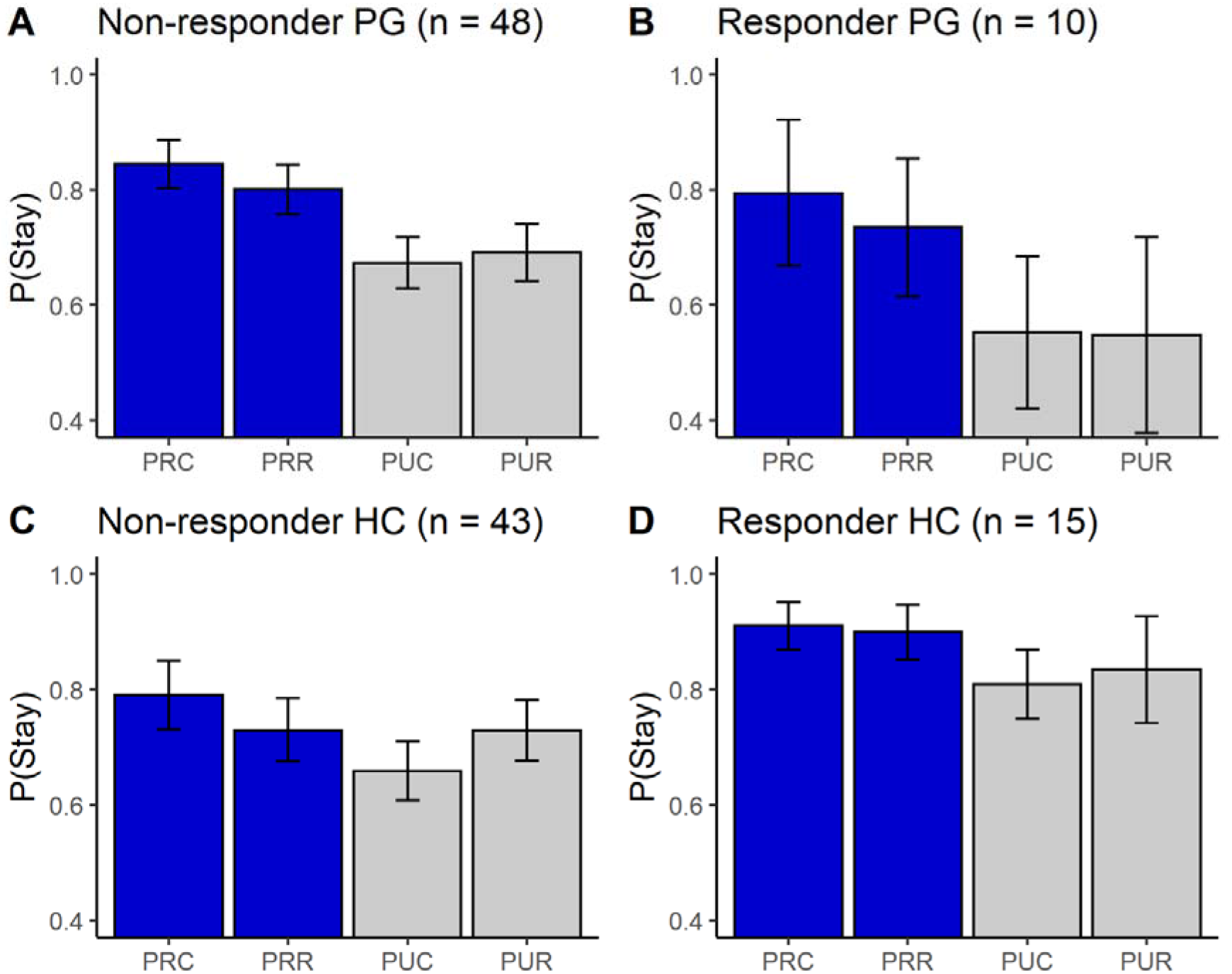
Probability to maintain a first step choice in pathological gamblers (PG) and healthy controls (HC) in trial following: a reward and a common transition (PRC), a reward and a rare transition (PRR), a loss and a common transition (PUC), as well as a loss and a rare transition (PUR). Graphs show mean values ± 2 SE.

A between-subject MANOVA (figure 5) was performed to evaluate the effects of diagnostic group (PG vs. HC), stress response (responders vs. non-responders), and their interaction on each computational parameter. Results indicated a significant interaction effect on the combined dependent variables (F(7, 106) = 4.23, p < .001; Wilks’ Λ = .78, η^2^_p_ = .22). Univariates analyses (displayed in supplementary table 1) indicated a significant effect of the interaction on the *ω*-parameter (F(1,112) = 7.79, p = .006, η^2^_p_ = .07). Simple effect analyses with Bonferroni corrections showed that the *ω*-parameter was significantly lower among responders HC than non-responders HC (F(1,56) = 12.90, p < .01, η^2^p = .19), but no significant difference was observed between responder and non-responder PG (F(1,56) = 0.22, p = .98, η^2^p = .004). The *ω*-parameter was significantly lower among PG than HC in the non-responder group (F(1,89) = 9.32, p = .01, η^2^p = .10), but not in the responder group (F(1,23) = 2.92, p = .34, η^2^p = .11). In six additional ANCOVA, we added the OSPAN, Raven, SCL90R, BDI, STAI-YA, and RS scores as covariables. The interaction effect remained significant (p < .05, see supplementary table 2).

**Figure 5.**
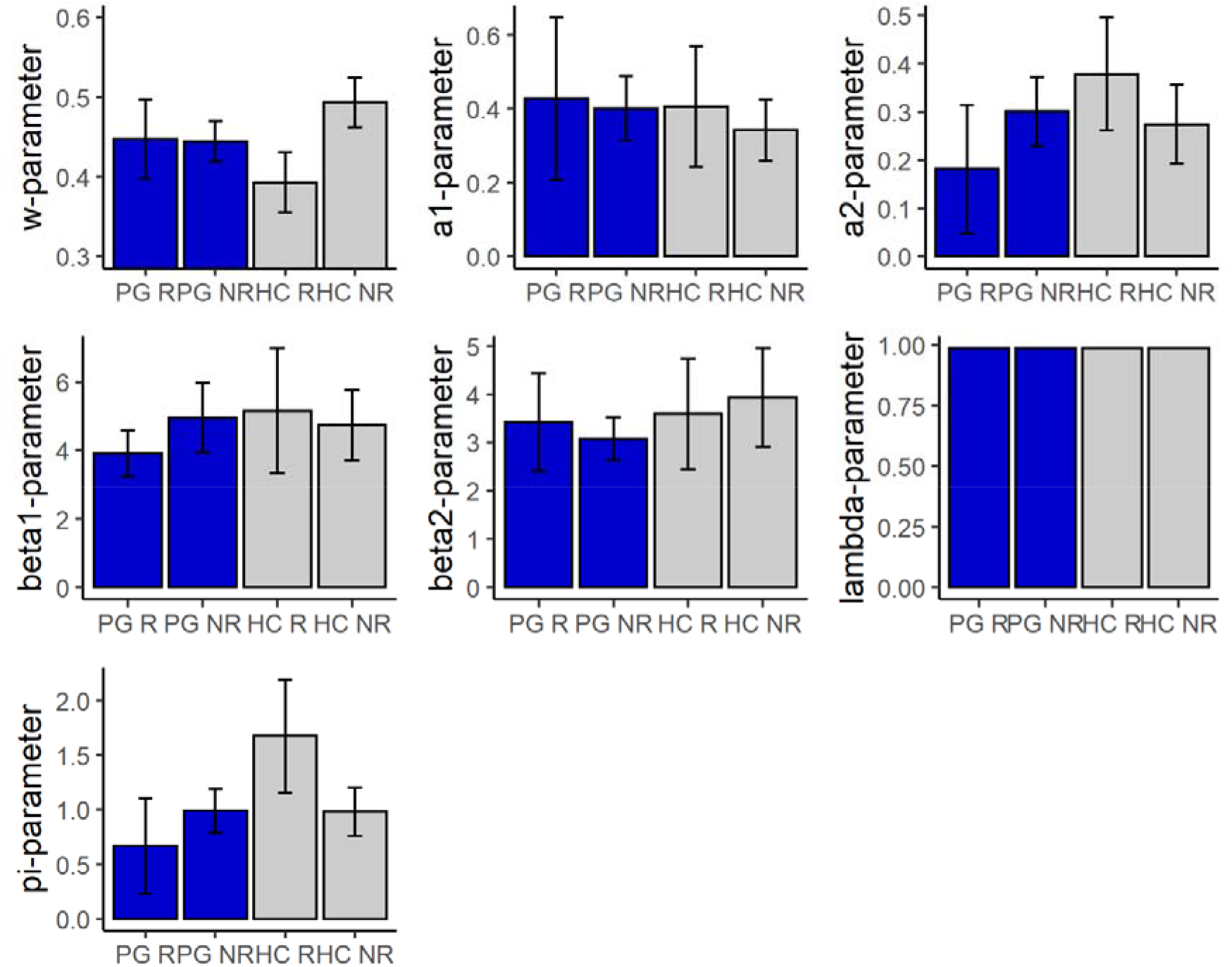
Mean of the computational parameters among responder pathological gamblers (PG S), non-responder pathological gamblers (PG NS), responder controls (HC S), and non-responder controls (HC NS). Graphs show mean values ± 2 SE.

Interestingly, the interaction also showed a significant effect on the ρ-parameter, (F(1,112) = 8.78, p = .004, η^2^_p_ = .07), an index quantifying perseveration. It was significantly lower among PG than HC in the stressed group (F(1,23) = 7.47, p = .047, η^2^_p_ = .25), but not in the non-responder group (F(1,89) = 0.003, p > .99, η^2^_p_ < .001). Additionally, the ρ-parameter was significantly higher among responder than non-responder HC (F(1,56) = 8.51, p = .02, η^2^_p_ = .13), but no significant difference was observed between responder and non-responder PG (F(1,56) = 1.70, p = .59, η^2^_p_ = .03).

The *ω*-parameter was regressed on the diagnostic group (DG; dummy coded: PG = −1 vs HC = 1), cortisol increase (dCort; continuous), and their interaction (figure 6). The model was significant (F(3,93) = 4.37, p < 0.01, R^2^ = 0.12), with a significant dCort*DG interaction (β = −0.56 (0.19), p = 0.004) indicating that stress was more deleterious on the *ω*-parameter among HC than PG. Bayesian Kendall’s correlations indicated strong evidence in favor of a negative correlation between the *ω*-parameter and dCort among HC (ρ(58) = −0.35, BF = 15.1), and moderate evidence against this correlation among PG (ρ(58) = −0.05, BF = 0.32). In six additional full factorial regressions, we added the OSPAN, Raven, SCL90R, BDI, STAI-YA, and RS scores as predictors. The interaction dCort*DG remained significant (p < 0.05) in each regression, and no additional predictor showed a significant main effect or interaction with DG and dCort (p > 0.05).

**Figure 1.**
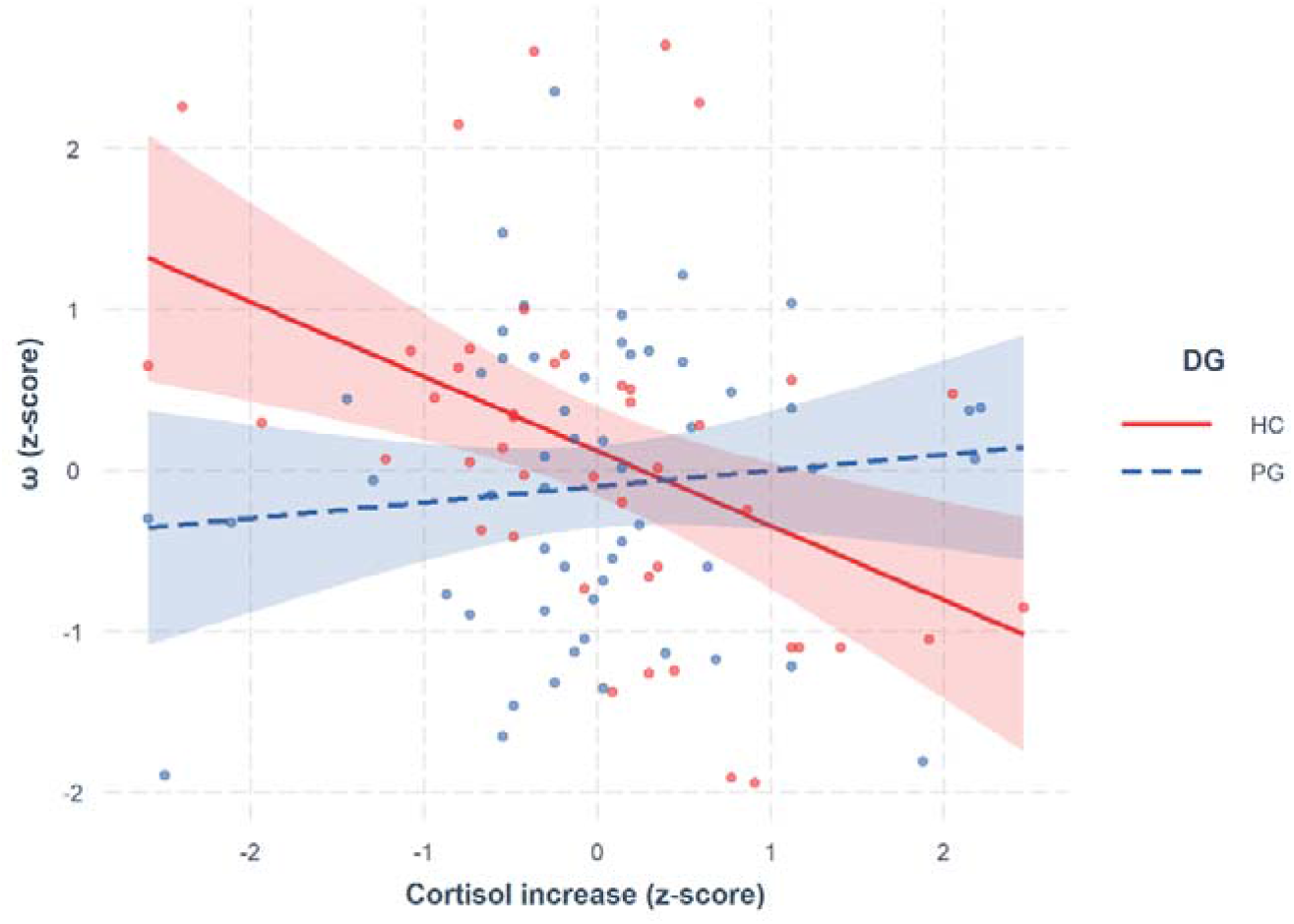
Effect of the cortisol increase on ω-parameter in pathological gamblers (PG) and healthy controls (HC), as determined by the computational model. Each continuous score was standardized, 95% IC is displayed around each regression line.

## DISCUSSION AND CONCLUSIONS

The objective of the present study was to investigate how MB and MF were orchestrated in GD according to a hybrid-RL 7-parameter computational model and whether PG were more sensitive than HC to the deleterious influence of stress on this dynamic. This is the first study to our knowledge that investigated the impact of stress on decision making in PG by employing a computational modelling approach that goes beyond the experimental approaches used in previous studies (e.g., Wyckmans et al., 2019).

Results indicated that the two groups had comparable baseline levels and stressor-induced elevation of salivary cortisol. A deleterious effect of enhanced cortisol level on MB/MF orchestration, characterized as a decrease of MB learning relative to MF learning, was found in HC but not in PG. Among non-stressed participants, results indicated that the balance between MB and MF contributions, formalized by the *ω*-parameter, was biased towards MF learning in PG. This pattern held even while controlling for the lower verbal working memory performance found in PG. To our knowledge, this is the first study to employ computational modelling to estimate these MF/MB parameters among PG. These results not only aligned with previous study with PG and MF/MB (Wyckmans et al., 2019) but also converged with similar studies conducted in subjects with substance-use disorders (Chen et al., 2021; Doñamayor et al., 2018; Sebold et al., 2017; Voon et al., 2015). Thus, these results support the idea that the inflexible repeated behaviors observed in PG might be underpinned by a greater inclination towards a habitual model of decision-making (Lucantonio et al., 2014). This tendency indicates a higher reliance towards MF-strategies, wherein choices are based essentially upon past rewards, with little consideration for changes in contingency between response and consequence.

Despite similar objective (cortisol) and subjective (ratings) responses to stress between the PG and HC groups, cortisol response was only associated with changes on the *ω*-parameter among HC (Otto, Raio, et al., 2013; Radenbach et al., 2015). Bayesian analyses further supported that the stress-induced cortisol increase was not associated with modulations in MB/MF orchestration in PG. A first explanation could be that the overall poorer working memory performance among PG and their general bias towards MF learning might explain the absence of a deleterious effect of the acute stressor on the MB/MF orchestration (i.e., floor effect). Indeed, evidence suggests that MB is partially supported by working memory (Culbreth et al., 2016), as it involves the learning of the action-outcome-transition contingencies and the planning of the following choices (Otto, Gershman, et al., 2013). Accordingly, the shift from MB towards MF learning among HC following stress might be the consequence of a stress-induced decrease in working memory performance (Wirz et al., 2018). Hence, the cortisol increase might have limited influence on PG’s already-impaired working memory, thereby restricting the modulation by stress on MB/MF orchestration in this population. However, group differences in the stress-induced modulation of learning strategies remained significant even when accounting for the effect of working memory performances, which suggests that this interpretation at best partially explain our results.

Another interpretation would be that the stress induced in the current study might have been insufficient among PG to elicit a behavioral adaptation during the Two-Step Markov Task. Despite similar cortisol increases across HC and PG, the latter are accustomed to functioning in highly stressful environments (Buchanan et al., 2020), which could be the cause of a persistent shift from goal-directed to habitual behavior (Dias-Ferreira et al., 2009). While we found no association between the *ω*-parameter and the social readjustment rating scale (SRRS; Holmes & Rahe, 1967), the latter does not account for stressful events specific to PG, such as financial preoccupation (Langham et al., 2015; Li et al., 2017), meaning that PG chronic stress might have been underestimated. Specific measures of chronic stress are therefore needed to further validate the influence of chronic stress on the behavioral response to acute stress in GD.

Additionally, interoception, which critically contributes to subjective emotional experience by modulating the subjective strength of bodily responses (i.e., the ability to perceive internal state) might have been abnormally inaccurate in PG, as a recent study suggests (Moccia et al., 2021). Also, in line with impaired interoception, alexithymic tendencies (e.g., difficulty in identifying feelings) are greater in PG than in healthy participants (Noël et al., 2018). This lack of accurate perception of internal bodily states related to emotions, such as the somatic experience of stress, might contribute to downplaying the influence of stress-induction on diverse behavioural responses among PG, including MF/MB orchestration. Of note, the visual analog scales used to score perceived stress was not meant to adequately capture interoceptive processing. A multidomain, multidimensional approach to interoception including cardiac, gastric, and respiratory assessments should be preferred in future studies to examine the role of this function in the association between acute stress and reinforcement learning in PG (Murphy et al., 2018; Wang et al., 2019).

Exploratory analyses showed that alongside the ω-parameter, the perseveration parameter (the maintenance of a choice irrespective of the previous outcomes and transitions) was higher among HC who showed a cortisol increase after the stress-inducing procedure. Stress-induced perseveration has never been reported and only one study found overall higher perseveration in binge-eating disorder and interpreted it as a lower involvement of cognitive control processes (Voon et al., 2015). While the ρ-parameter did not significantly differ between HC and PG, exacerbated perseveration among HC following acute stress might result from the deleterious effect of cortisol increase on the prefrontal cortex (Wirz et al., 2018).

As computational modelling allows us to quantify inter-individual differences in mechanisms underlying addiction development and maintenance, they hold great promises to promote targeted clinical interventions (Heinz et al., 2017). For instance, contingency management therapies, which show a high level of efficacy in substance use disorders (Davis et al., 2016; Mantzari et al., 2015), might specifically help PG displaying low MB capacities, as MB strategy use is increased when higher incentives are offered (Kool et al., 2017; Patzelt et al., 2019). Episodic future thinking training represents another promising way to reduce addiction severity (Snider et al., 2016). As outlined, PG encounter difficulties in mentally simulating future events in a vivid manner (Noël et al., 2017). Episodic foresight might support model-based control in the sense that the agent prospectively evaluates actions based on their potential outcomes. However, little is known about the relationship between learning an internal model to prospectively make decisions and the capacity to mentally navigate the future. Finally, several brain stimulation techniques (tDCS, rTMS) might target critical neural networks involved in MB/MF orchestration, thus representing a promising way to treat gambling addiction (Smittenaar et al., 2013; Weissengruber et al., 2019).

The present study is not exempt from limitations. First, only a third of participants undergoing the SECPT showed a cortisol increase, lowering our power to observe small cortisol-induced behavioral changes. Nevertheless, both the HC and the PG groups displayed similar levels of cortisol stress response, and the influence of acute stress on HC’s decisional processes was consistent with the literature (Otto, Raio, et al., 2013; Radenbach et al., 2015). Additionally, false beliefs about action and reward contingencies prevalent in GD, such as the hot-hand effect or gambler’s fallacy (Joukhador et al., 2004), have not been taken into account. As MB and MF are high-level processes, emerging from many sub-computations (Collins & Cockburn, 2020; Daw, 2018), these alternative decision strategies could have severely biased our strictly dichotomized behavioral assessment (Mohr et al., 2018). Further iterations of this paradigm in GD should therefore control for these false beliefs and assess how they might be related to weakened MB learning.

To summarize, the present study adds evidence towards the hypothesis that GD includes a compulsive component with a propensity towards inflexible habits formation when facing new decisions, arising from a bias towards MF control. PG’s resistance to acute stress might be caused by several non-exclusive factors, namely low working-memory capacities, high chronic stress, and deleterious interoceptive accuracy. Despite a normal cortisol response to stress, we found an abnormal pattern of the modulation of stress on the relative weighing of goal-directed and habitual action strategies in GD.

