## Supplementary Material for "The modulation of acute stress on Model-Free and Model-Based reinforcement learning in Gambling Disorder"

### Method

#### Salivary cortisol assessment

Patients received tubes and cotton swabs to collect saliva samples. Participants were asked to place a cotton swab under their tongue for 2 min. Participants were instructed to refrain from smoking, eating, or brushing their teeth for at least 30 min before, and consuming alcohol for 12 h before cortisol collection. Cortisol levels were determined using a commercially available competitive enzyme linked immunosorbent assay (ELISA) (Salimetrics® LLC, Carlsbad, CA, USA). Samples were thawed, vortexed and centrifuged at 1500×g for 15 min and clear samples pipetted into appropriate wells. Cortisol in standards and samples compete with cortisol conjugated to horseradish peroxidase for the antibody binding sites on a microtiter plate. After incubation, unbound components are washed away. Bound cortisol enzyme conjugate is measured by the reaction of the horseradish peroxidase enzyme to the substrate tetramethylbenzidine (TMB). This reaction produces a blue color. A yellow color is formed after stopping the reaction with an acidic solution. The optical density is read on a standard plate reader at 450 nm. The amount of cortisol enzyme conjugate detected is inversely proportional to the amount of cortisol present, in the sample.

#### Computational modeling

The outcome of each second-step stages, the transition type and the response time were recorded for each trial. Data were then fitted on a 7-parameter model as proposed by Daw et al. (2011). The RL task consists of three states: the first step stage $s_{A}$ and both second step stages $s_{B}$ and $s_{C}$. In each state, participant has the choice between two actions: $a_{A}$ and $a_{B}$. The transition function $T\left( s, a, s^{'} \right)$ dictates state $s^{'}$ in which the agent will transition into if action $a$ is performed while in state $s$.

$$T\left( s, a, s^{'} \right)=P(s_{i+1}=s^{'}\left| s_{i}=s, a_{i}=a) \right.$$

By making choice, the agent go from state $s_{i}$ to state $s_{i+1}$to maximize his reward. Since the rewards is stochastically dependent on the state t, the reward function is defined as the average reward in that state.

$$R\left( s \right)= E\left[ \left. r_{i} \right|s_{i}=s \right]$$

##### Model-Free RL algorithm

The algorithm accounting for habitual learning is the Model-Free State-Action-Reward-State-Action (SARSA) (λ) temporal difference learning algorithm (TD; Rummery & Niranjan, 1994). It predicts the value of each state-action pairs and updates these over the learning according to:

$$Q_{MF}\left( s_{i,t+1},a_{i,t+1} \right)= Q_{MF}\left( s_{i,t},a_{i,t} \right)+ \alpha_{i}\delta_{i,t}$$

The teaching signal $\delta_{i,t}$ is the reward prediction error and corresponds to the difference between the expected and the actual reward:

$$\delta_{i,t}= r_{i,t}+ Q_{MF}\left( s_{i,t+1},a_{i,t+1} \right)- Q_{MF}\left( s_{i,t},a_{i,t} \right)$$

Since there are no third step in the RL task, $Q_{MF}\left( s_{3,t},a_{3,t} \right)$ is always equal to 0. Besides, no reward is delivered after the first step and $r_{1,t}$ is therefore always equal to 0 too. To assess the impact of the second-step reward on the first-step choice, the eligibility parameter λ is introduced to propagate the second step $Q_{MF}\left( s_{2,t},a_{2,t} \right)$ towards the next first step $Q_{MF}\left( s_{1,t},a_{1,t} \right)$.

$$Q_{MF}\left( s_{1,t+1},a_{1,t+1} \right)= Q_{MF}\left( s_{1,t},a_{1,t} \right)+ \alpha_{1}\lambda\delta_{2,t}$$

This eligibility parameter reflects the main effect of the reward, and the TD algorithm therefore captures habitual learning as it updates choice values only retrospectively without taking the goal-representation into account. Importantly, it does not immediately map on habits, which are divorced from their consequences (Reiter et al., 2016).

##### Model-Based RL algorithm

Since the TD does not take the transitions into account, a second algorithm, the model-based RL algorithm, is used to fit the data. It takes the task structure into account by including the transition matrix. It maps each first-stage action to second-state stages and multiplies it by the maximum value at the second stage. Since there are no transition in the second step, $Q_{MB}\left( s_{2,t},a_{2,t} \right)= Q_{MF}\left( s_{2,t},a_{2,t} \right)$.

$$Q_{MB}\left( s_{A},a_{j} \right)=P\left( s_{B} | s_{A},a_{j} \right) \max_{a\in\left\{ a_{A},a_{B} \right\}} Q_{MF}\left( s_{B},a \right)+ P\left( s_{C} | s_{A},a_{j} \right) \max_{a\in\left\{ a_{A},a_{B} \right\}} Q_{MF}\left( s_{C},a \right)$$

##### Hybrid model

The hybrid algorithm connects the MF and the MB algorithms via a weighing parameter ω. It captures the relative influence of the MB and MF algorithms where higher values (0.5 < ω < 1) show a higher reliance upon MB computations, and lower values (0 < ω < 0.5) to a higher reliance upon MF computations.

$$Q_{net}\left( s_{A},a_{j} \right)= \omega Q_{MB}\left( s_{A},a_{j} \right)+(1-\omega)Q_{TD}\left( s_{A},a_{j} \right)$$

The action probability is then computed with a SoftMax function on $Q_{net}.$

$$P\left( a_{i, t}=a | s_{i,t} \right)= \frac{exp(\beta_{i}\left[ Q_{net}\left( s_{i,t},a \right)+p \cdot rep(a) \right]}{\sum_{a'} exp(\beta_{i}\left[ Q_{net}\left( s_{i,t},a' \right)+p \cdot rep(a') \right]}$$

Seven parameters are therefore computed with the hybrid model and can then be compare between-, within-subjects:

- The weighing parameter ω which captures the balance between MB and MF reliance.
- The inverse temperature β_1_ and β_2_ correspond to the stochasticity of each step choices with higher value reflecting higher influence of the model on the choice. It is separately computed for each choice to be consistent with the different levels of stochasticity of each step.
- The free parameters α_1_ and α_2_ corresponds to the free learning-rate parameter in each step.
- The free parameter ρ captures perseveration during the first step with $rep\left( a \right)$ reflecting the repetition of the previous first step action ($rep\left( a \right)=1$) or switching ($rep\left( a \right)=0$).
- And finally, the eligibility parameter λ reflects the main effect of the reward on subsequent choice.

#### Working memory assessment

The operation span (OSPAN) was used to assess working memory performances (Turner & Engle, 1989; Unsworth et al., 2005). It is composed of 15 blocks. Each one is composed of 2 to 6 solved arithmetical operations (3 blocks of 2 operations, 3 blocks of 3 operations, 3 blocks of 4 operations, 3 blocks of 5 operations and 3 blocks of 6 operations). The participants must tell if the operation’s answer is right or wrong by pressing one of two keys on a dedicated keyboard (green for right and red for wrong). Words appear between each operation, and it is asked to the participants to remember them. After each block, they must return the words in the same order by typing them. The number of words to remember therefore depends on the number of operations. The Partial Credit Unit technique (Conway et al., 2005) was used to estimate a working memory score (mean proportion of words correctly given at the end of each block).

Besides, each participant’s fluid intelligence was assessed through the nine-item forms of the Raven’s Standard Progressive Matrices Test (RSPM). It consists in 9 multiple-choice pattern matching tasks where participant must identify the piece required to complete a sequence of pattern (Raven, 2000). The items we used were selected by Bilker et al. (2012) from the 60 original items as they offer similar psychometric characteristics and 75% administration time savings compared to the 30-item version.

#### Clinical questionnaires

Alcohol use disorder was assessed by the Alcohol-Use Disorder Identification Test (Saunders et al., 1993), nicotine consumption by the Fagerstöm Test for Nicotine Dependence (Heatherton et al., 1991), psychiatric symptoms and comorbidities by the SCL-90R (Derogatis & Cleary, 1977), negative affect by the Beck Depression Inventory (Beck et al., 1988) and the PANAS scale (Watson et al., 1988), reward and punishment sensitivity by the SPSRQ (Lardi et al., 2008), impulsivity by the UPPS (Van der Linden et al., 2006), state and trait anxiety by the STAI-YA and STAI-YB (Bruchon-Schweitzer & Paulhan, 1993), as well as habit reliance with the Creature Of Habit Scale (Wyckmans et al., 2020)

### Supplementary analyses

##### Analyses of variance

A between-subject MANOVA (figure 5) was performed to evaluate the effects of diagnostic group (PG vs. HC), stress response (responders vs. non-responders), and their interaction on each computational parameter. Results indicated a significant interaction effect on the combined dependent variables (F(7, 106) = 4.23, p < .001; Wilks' Λ = .78, η^2^_p_ = .22).

| Parameter | Effects | F(1,112) | p | η^2^_p_ |
| --- | --- | --- | --- | --- |
| α1 | DG | 0.44 | .51 | .004 |
|  | SG | 0.69 | .41 | .01 |
|  | SG*DG | 0.09 | .76 | .001 |
| β1 | DG | 0.27 | .61 | .002 |
|  | SG | 0.06 | .80 | .001 |
|  | SG*DG | 0.76 | .38 | .01 |
| α2 | DG | 2.71 | .10 | .02 |
|  | SG | 0.009 | .92 | < .001 |
|  | SG*DG | 3.77 | .06 | .03 |
| β2 | DG | 0.35 | .56 | .003 |
|  | SG | 0.04 | .84 | < .001 |
|  | SG*DG | 0.07 | .79 | .001 |
| ρ | **DG** | **9.04** | **.003** | **.08** |
|  | SG | 1.38 | .24 | .01 |
|  | **SG*DG** | **8.77** | **.004** | **.07** |
| *ω* | DG | .005 | .94 | < .001 |
|  | **SG** | **4.51** | **.04** | **.04** |
|  | **SG*DG** | **7.79** | **.01** | **.07** |
| λ | DG | 0.32 | .57 | .003 |
|  | SG | 1.68 | .20 | .02 |
|  | SG*DG | 0.39 | .54 | .003 |
| Supplementary table 1. Results of the univariate analyses assessing the effect of the diagnostic group (DG), the stress group (SG), and their interaction on each computational parameter. Significant results are displayed in bold. | | | | |

##### Analyses of covariance

ANCOVA were performed to assess the effect of the stress group, the diagnostic group, and their interaction on the *w*-parameter, while accounting for the main effect of the covariate, as well as its two-way interactions with the stress and the diagnostic groups.

As HC and PG differed on working memory performance (OSPAN scores; U(116) = 1207, p = .009), and WM capacities being reported to moderate the deleterious stress effect on MB learning (Otto, Raio, et al., 2013), we assessed whether differential OSPAN scores would mediate the interaction between SG and DG on the *ω*-parameter. An ANCOVA was performed to evaluate the effect of the diagnostic group (DG; PG vs HC), the stress group (SG: stressed vs not-stressed), the OSPAN score (continuous), and each two-way interaction on the *ω*-parameter. Results showed a significant main effect of SG (F(1,109) = 7.48, p = .007; η^2^_p_ = .06) and of the interaction between SG and OSPAN (F(1,109) = 10.26, p = .002; η^2^_p_ = .09), while the interaction between SG and DG remained significant (F(1,109) = 5.71, p = .02; η^2^_p_ = .05). It indicated that while the WM capacities indeed moderated the stress effect on the MB/MF balance, it did not mediate the effect of the diagnostic group. All other effects where non-significant (p_s_ > .05). Other covariates, that is, the Raven score, state-anxiety, depression symptoms, psychiatric comorbidities, and reward sensitivity were included in the model in the same way. None showed a significant main or moderation effect (p_s_ > .05). Full results are displayed in supplementary table 2.

|  | **OSPAN** | | |  | **BDI** | | |
| --- | --- | --- | --- | --- | --- | --- | --- |
|  | F (1, 109) | p | η^2^_p_ |  | F (1, 109) | p | η^2^_p_ |
| DG | 3.8 | .05 | .03 | DG | 0.11 | .74 | .001 |
| **SG** | **7.48** | **.01** | **.06** | SG | 1.79 | .18 | .02 |
| OSPAN | 0.62 | .44 | .01 | BDI | 0.15 | .70 | .001 |
| **DG*SG** | **10.26** | **.002** | **.09** | **DG*SG** | **7.66** | **.007** | **.07** |
| SG*OSPAN | 3.9 | .05 | .04 | SG*BDI | 0.05 | .82 | < .001 |
| **DG*OSPAN** | **5.71** | **.02** | **.05** | DG*BDI | 0.09 | .76 | .001 |
|  | **Raven** | | |  | **STAI-YA** | | |
|  | F (1, 109) | p | η^2^_p_ |  | F (1, 109) | p | η^2^_p_ |
| DG | 0.04 | .84 | < .001 | DG | 0.66 | .42 | .006 |
| SG | 0.04 | .84 | < .001 | SG | 0.2 | .66 | .002 |
| Raven | 1.37 | .25 | .01 | STAIYA | 0.37 | .54 | .003 |
| **DG*SG** | **6.51** | **.01** | **.06** | **DG*SG** | **9.18** | **.003** | **.08** |
| SG*Raven | 0.15 | .70 | .001 | SG*STAIYA | 0.75 | .39 | .007 |
| DG*Raven | 0.01 | .94 | < .001 | DG*STAIYA | 0.77 | .38 | .007 |
|  | **SCL-90R** | | |  | **RS** | | |
|  | F (1, 109) | p | η^2^_p_ |  | F (1, 109) | p | η^2^_p_ |
| DG | 0.09 | .76 | .001 | DG | 0.002 | .97 | < .001 |
| SG | 0.88 | .35 | .008 | SG | 2.13 | .15 | .02 |
| SCL90R | 0.001 | .98 | < .001 | RS | 0.84 | .36 | .01 |
| **DG*SG** | **7.75** | **.01** | **.07** | **DG*SG** | **4.78** | **.03** | **.04** |
| SG*SCL90R | 0.15 | .70 | .001 | SG*RS | 0.99 | .32 | .01 |
| DG*SCL90R | 0.17 | .68 | .002 | DG*RS | 0.007 | .93 | < .001 |
| Supplementary table 2. Results of the six ANCOVAs assessing the effect of the stress group (SG), the diagnostic group (PG), and their interaction on the w-parameter, while controlling respectively for the OSPAN score, the Raven score, the psychiatric comorbidities (SCL90R), the depressive symptoms (BDI), the state-anxiety (STAIYA), and the reward sensitivity (RS). Significant results are displayed in bold. | | | | | | | |

##### Correlation analyses

Pearson’s correlations were performed between each computational parameter and each clinical variable. To avoid the confounding effect of the induced stress, they were performed separately for the stressed and the not-stressed group. Results are displayed in supplementary tables 3 and 4.

| Responders | a1 | b1 | a2 | b2 | pi | w | lambda |
| --- | --- | --- | --- | --- | --- | --- | --- |
| AUDIT | -0.04 | 0.34 | -0.10 | 0.06 | -0.24 | -0.10 | -0.07 |
| SOGS | 0.19 | -0.20 | -0.48 | -0.09 | **-0.62**** | 0.35 | 0.05 |
| DSM | 0.11 | -0.13 | -0.44 | -0.19 | **-0.48*** | 0.37 | -0.07 |
| OSPAN | 0.35 | **-0.50*** | 0.32 | 0.13 | -0.12 | -0.36 | 0.26 |
| Raven | 0.13 | -0.14 | **0.42*** | 0.05 | 0.14 | -0.08 | 0.12 |
| SCL90R | -0.07 | -0.09 | -0.05 | 0.11 | -0.24 | 0.07 | 0.09 |
| BDI | -0.27 | 0.12 | -0.26 | **0.40*** | -0.14 | 0.11 | -0.14 |
| PANAS pos | -0.09 | -0.25 | **0.42*** | **-0.4*** | 0.25 | 0.19 | -0.29 |
| PANAS neg | -0.33 | -0.09 | 0.01 | 0.12 | 0.002 | 0.07 | -0.31 |
| STAI-YA | 0.15 | -0.11 | -0.25 | 0.10 | -0.39 | -0.03 | 0.25 |
| STAI-YB | -0.16 | 0.02 | -0.15 | 0.20 | -0.21 | -0.05 | 0.05 |
| SRRS | -0.12 | -0.28 | 0.18 | 0.02 | -0.11 | -0.03 | -0.13 |
| PS | 0.06 | -0.06 | -0.14 | 0.27 | -0.11 | -0.10 | 0.15 |
| RS | -0.04 | -0.22 | -0.03 | 0.15 | -0.11 | 0.33 | -0.13 |
| CoH R | -0.29 | 0.06 | -0.28 | 0.03 | 0.02 | 0.26 | 0.06 |
| CoH A | 0.12 | 0.12 | -0.003 | -0.18 | -0.30 | 0.10 | 0.49 |
| NU | -0.14 | -0.17 | -0.03 | -0.19 | -0.29 | 0.25 | -0.19 |
| PU | **-0.45*** | 0.11 | 0.01 | -0.04 | -0.02 | 0.19 | -0.28 |
| LoPr | -0.16 | 0.09 | -0.31 | 0.32 | -0.02 | 0.11 | -0.32 |
| LoPe | 0.06 | -0.004 | **-0.48*** | 0.32 | -0.15 | -0.16 | 0.04 |
| SS | -0.02 | -0.06 | 0.11 | -0.35 | -0.16 | 0.19 | <0.001 |
| Supplementary table 3. Pearson's coefficient resulting from the correlation among responders between each computational parameter and alcohol use disorder symptoms (AUDIT), gambling disorder symptoms (SOGS), number of DSM items (DSM), WM capacities (OSPAN), Raven score, psychiatric comorbidities (SCL90R), depressive symptoms (BDI), positive affects (PANAS pos), negative affects (PANAS neg), state-anxiety (STAI-YA), trait-anxiety (STAI-YB), chronic stress (SRRS), sensitivity to punition (PS), sensitivity to reward (RS), routine tendencies (CoH R), automatism tendencies (CoH A), negative urgency (NU), positive urgency (PU), lack of premeditation (LoPr), lack of perseverance (LoPe), and sensation seeking (SS). Significant results are displayed in bold. * p < .05; **p < .001 | | | | | | | |

| Non-Responders | a1 | b1 | a2 | b2 | pi | w | lambda |
| --- | --- | --- | --- | --- | --- | --- | --- |
| AUDIT | 0.11 | -0.09 | 0.01 | 0.06 | 0.03 | 0.04 | -0.02 |
| SOGS | 0.02 | 0.00 | 0.17 | -0.12 | 0.15 | -0.03 | 0.01 |
| DSM | 0.04 | -0.01 | 0.01 | -0.17 | -0.02 | 0.05 | -0.06 |
| OSPAN | -0.06 | -0.05 | **0.33**** | -0.13 | 0.19 | **0.28**** | -0.03 |
| Raven | 0.11 | 0.02 | **0.36**** | -0.14 | 0.12 | **0.30**** | 0.11 |
| SCL90R | 0.02 | 0.06 | 0.06 | -0.10 | 0.06 | 0.01 | 0.18 |
| BDI | -0.04 | 0.05 | 0.08 | -0.09 | 0.06 | 0.01 | 0.17 |
| PANAS pos | 0.03 | 0.04 | -0.05 | 0.03 | -0.09 | 0.05 | -0.03 |
| PANAS neg | -0.03 | 0.02 | -0.01 | -0.06 | 0.04 | 0.02 | 0.13 |
| STAI-YA | 0.03 | 0.13 | -0.03 | 0.08 | 0.01 | -0.03 | 0.17 |
| STAI-YB | -0.08 | 0.15 | 0.04 | 0.09 | 0.11 | 0.03 | 0.15 |
| SRRS | 0.01 | -0.02 | 0.07 | -0.04 | 0.13 | 0.12 | 0.07 |
| PS | -0.12 | 0.11 | 0.01 | -0.01 | -0.05 | -0.06 | -0.03 |
| RS | -0.11 | -0.06 | 0.10 | **-0.23*** | 0.02 | -0.08 | -0.04 |
| CoH R | -0.08 | -0.09 | 0.21 | -0.31 | 0.10 | -0.30 | 0.12 |
| CoH A | 0.07 | -0.12 | 0.08 | -0.08 | 0.12 | -0.15 | **0.39*** |
| NU | -0.12 | -0.03 | **-0.24*** | -0.03 | -0.05 | 0.00 | **-0.22*** |
| PU | -0.10 | -0.01 | -0.04 | 0.05 | -0.04 | -0.02 | -0.12 |
| LoPr | 0.06 | -0.03 | -0.10 | -0.13 | -0.06 | -0.08 | 0.00 |
| LoPe | 0.00 | -0.01 | -0.07 | -0.08 | -0.06 | 0.05 | 0.01 |
| SS | -0.16 | -0.01 | 0.11 | -0.11 | 0.10 | 0.12 | -0.08 |
| Supplementary table 4. Pearson's coefficient resulting from the correlation among non-responders between each computational parameter and alcohol use disorder symptoms (AUDIT), gambling disorder symptoms (SOGS), number of DSM items (DSM), WM capacities (OSPAN), Raven score, psychiatric comorbidities (SCL90R), depressive symptoms (BDI), positive affects (PANAS pos), negative affects (PANAS neg), state-anxiety (STAI-YA), trait-anxiety (STAI-YB), chronic stress (SRRS), sensitivity to punition (PS), sensitivity to reward (RS), routine tendencies (CoH R), automatism tendencies (CoH A), negative urgency (NU), positive urgency (PU), lack of premeditation (LoPr), lack of perseverance (LoPe), and sensation seeking (SS). Significant results are displayed in bold. * p < .05; **p < .001 | | | | | | | |

##### Response time

Response times at each step were analyzed with repeated measures ANOVA. We expected the first step choice to be faster after an unrewarded trial, especially among PG. Besides, MB learning taking the transition into account, we expected the second step choice to be slower after a rare transition when the decisional balance weights towards MB learning (Wyckmans et al., 2019).

A repeated measures ANOVA was performed to evaluate the effects of the diagnostic group (DG: PG vs HC), the stress group (SG: stressed vs not-stressed), the outcome of the previous trial (within-subject, rewarded vs non-rewarded), and their interaction on the response time during the first step. Results show only a significant effect of the reward (F(1,112) = 16.64, p < .001; η^2^_p_ = .13), indicating that participants made a choice faster after an unrewarded (mean ± sd RT = 391.58 ± 100.85 ms) than a rewarded trial (mean ± sd RT = 412.08 ± 98.64 ms).

A repeated measures ANOVA was performed to evaluate the effects of the diagnostic group (DG: PG vs HC), the stress group (SG: stressed vs not-stressed), the transition type (within-subject, common vs rare), and their interaction on the response time during the second step. Results show only a significant effect of the transition (F(1,112) = 47.90, p < .001; η^2^_p_ = .30), indicating that participants made their choice slower after a rare (mean ± sd RT = 619.90 ± 182.53 ms) than after a common transition (mean ± sd RT = 559.05 ± 149.36 ms).

Deltas (dRT1 and dRT2) were computed by subtracting the response time of choices following a rewarded trial/common transition, to the response time of choices following an unrewarding trial/rare transition, respectively. Pearson’s correlations were performed between the computational parameters and the deltas. Their results are indicated in supplementary table 5. Crucially, the *ω-*parameter was significantly and positively associated with dRT2, indicating that the more participants slowed their response time after a rare transition, the more the MB/MF balance would weight towards MB learning.

|  | **dRT1** | **dRT2** |
| --- | --- | --- |
| α1 | -0.16 | -0.09 |
| β1 | -0.01 | 0.09 |
| α2 | 0.03 | 0.36*** |
| β2 | 0.03 | -0.01 |
| ρ | 0.05 | 0.19* |
| ω | 0.02 | 0.30*** |
| λ | -0.20 | -0.08 |
| Supplementary table 5 Coefficient of the Pearson's correlation between the computational parameters and the differences of reaction time: during the first choice following a rewarded or unrewarded outcome (dRT1), as well as during the second choice following a common or rare transition (dRT2). * p < .05, ** p < .01, *** p < .001 | | |
